# Norepinephrine stimulates protein synthesis in astrocytes

**DOI:** 10.1101/2025.07.27.665649

**Authors:** Florencia Cabrera-Cabrera, Helena Tull, Giulia Urone, Age Utt, Robert Zorec, Indrek Koppel

## Abstract

Norepinephrine is a neuromodulator that can activate multiple subcellular signalling pathways to regulate various physiological responses in different cell types. In astrocytes, it is known to have a wide range of metabolic effects, including regulation of glucose uptake and glycolysis, glycogen breakdown and lactate production. Here we report that norepinephrine stimulates protein synthesis in rat primary cortical astrocytes and the C6 glioma cell line. The mTOR-pS6K pathway is engaged during the norepinephrine-mediated induction of protein synthesis, which is abolished by mTOR inhibition and appears to be mediated mainly by β-adrenergic receptors. We show that blocking glycolysis with 2-deoxy-D-glucose inhibits both basal and norepinephrine-induced translation in astrocytes. Finally, we demonstrate that the norepinephrine-mediated increase in protein synthesis is preceded by a decrease in astrocytic ATP levels. Our results reveal a novel regulatory role for norepinephrine in the central nervous system, tightly controlling protein synthesis in astrocytes.

## Introduction

Norepinephrine is a neuromodulator that regulates multiple physiological processes in different central nervous system (CNS) cell types (O’Donnell et al., 2012). Released from axonal varicosities of *locus coeruleus* neurons, norepinephrine activates its cognate G protein-coupled receptors (GPCRs) expressed in both neurons and glia (O’Donnell et al., 2012). Astrocytes express many adrenergic receptors, including α_1_, α_2_, β_1_ β_2_ and β_3_ receptors (Hertz et al., 2010; Zhang et al., 2014; Koppel et al., 2018; D’Adamo et al., 2021), allowing flexible norepinephrine concentration-dependent modulation of cAMP- and Ca^2+^-dependent signalling pathways (Wahis and Holt, 2021). In astrocytes, norepinephrine regulates glucose uptake and glycolysis, glycogen breakdown and lactate production (Bak et al., 2018; Fink et al., 2021; Zorec and Vardjan, 2023), which has been associated with powering long-lasting molecular changes in neurons that underlie learning and memory (Alberini et al., 2018). Other known adrenergic effects on astrocytes include enhancement of glutamate uptake by α_1_ receptor activation (Alexander et al., 1997) and increased extracellular K^+^ clearance by norepinephrine (Hajek et al., 1996). At the level of synaptic modulation, activation of astrocytic adrenoreceptors has recently been shown to play a key mediating role in synaptic plasticity in several animal models (Chen et al., 2025; Guttenplan et al., 2025; Lefton et al., 2025).

Protein synthesis is a core cellular anabolic process that needs to be tightly regulated (Livingstone et al., 2010). Protein synthesis regulation is mediated through different intracellular signalling pathways, which integrate information about cellular energy levels, amino acid availability and other parameters (Livingstone et al., 2010). Growth factors are well-known agents that rapidly promote protein synthesis via PI3K/Akt/mTOR and MAPK signalling pathways (Saxton and Sabatini, 2017; Roux and Topisirovic, 2018). GPCR-mediated regulation of protein synthesis has been studied less thoroughly (Tréfier et al., 2018), although the effect of class I mGluR agonist DHPG on increasing neuronal protein synthesis is well established (Huber et al., 2000; Raymond et al., 2000).

Norepinephrine has been shown to stimulate protein synthesis in cardiomyocyte cultures (Simpson, 1985; Dubus et al., 1990). Both α_1_ and β adrenergic receptors were implicated in cardiac hypertrophy and protein synthesis stimulation (Simpson, 1985; Dubus et al., 1990; Decker et al., 1993; Pinson et al., 1993). It is not known, however, if norepinephrine can exert a similar protein synthesis-enhancing effect in the cells of the nervous system. Here we show that norepinephrine induces protein synthesis in cultured rat cortical astrocytes and C6 glioma cells, suggesting a novel role for this common neuromodulator in the CNS.

## Results

### Norepinephrine stimulates protein synthesis in primary cultured astrocytes

Given the metabolic and morphological effects reported for norepinephrine (NE) in astrocytes, we wondered how it would affect the rate of protein synthesis. To test this, we treated serum-starved primary cortical astrocytes with NE for up to 3 h, followed by pulse labelling with puromycin (PMY) to detect newly synthesized proteins (Schmidt et al., 2009). We then quantified puromycinylated proteins by Western Blot and observed that NE elicits an increase in protein synthesis over time, similar to that induced by the addition of foetal bovine serum (Fig. 1A, B). Measurement of ^35^S-Met/Cys incorporation confirmed this effect (Fig. 1C). In addition, we used PMY immunocytochemistry analysis, detecting increased incorporation of PMY into nascent proteins in astrocytes treated with NE for 3h (Fig. 1D, E). A similar effect was detected in the C6 glioma cell line (Fig. S1A), and to a lesser extent in primary cortical neurons (Fig. S1B), but not in the HMC3 microglial cell line (Fig. S1C), suggesting that a NE-mediated increase in protein synthesis manifests mainly in astrocytes.

**Figure 1.**
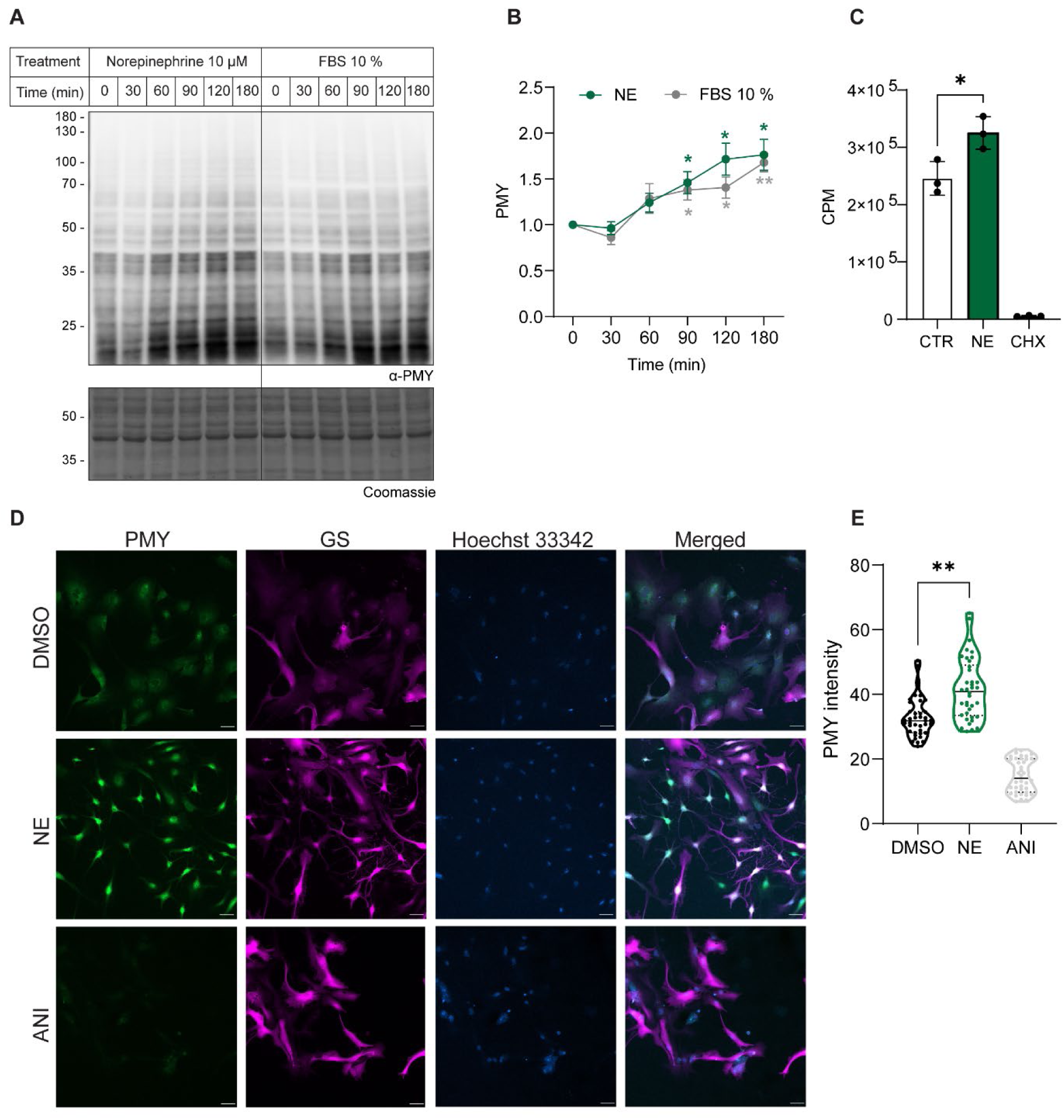
Norepinephrine stimulates protein synthesis in cultured cortical astrocytes. **A.** Serum-deprived primary cultures of rat cortical astrocytes were treated with 10 µM norepinephrine (NE) or 10% foetal bovine serum (FBS) for the indicated periods, newly synthesised proteins were labelled with 5 µM puromycin (PMY) during the final 15 minutes of treatment and detected by Western Blot using an anti-PMY antibody. Membranes were stained with Coomassie as a loading control. **B.** Quantification of signal intensity normalised to DMSO controls is shown as mean ± SEM for n=5 biological replicates. *p < 0.05; ** p < 0.01 (one sample t-test comparing the means with a control mean of 1). **C.** Incorporation of ^35^S-Met/Cys into *de novo* synthesised proteins in serum-deprived astrocytes treated with 10 µM NE for 3 h (or DMSO, CTR). As a control of protein synthesis inhibition, cells were co-treated with 100 µg/mL cycloheximide (CHX). Shown here are counts per minute (CPM) for n=3 biological replicates (mean ± SEM, *p < 0.05, one-way ANOVA with a Tukey’s multiple comparison test). **D.** Newly synthesised proteins from NE- or DMSO-treated astrocytes for 3 h were labelled with 5 µM PMY for 2.5 min and analysed by immunocytochemistry using an anti-PMY antibody (green). Staining with anti-GS antibody was included as an astrocytic marker (magenta). Cells were treated with the protein synthesis inhibitor anisomycin (10 µg/mL, ANI) as a negative signal control. Scale bar: 50 µm. **E.** PMY signal intensity on GS-positive cells was quantified using ImageJ from 10 different fields for each biological replicate (n=4). ** p < 0.01 (Kruskal-Wallis test with Dunn’s correction for multiple comparisons).

### The mTOR pathway is engaged during the NE-mediated induction of protein synthesis

To further characterize the effect of NE on global protein synthesis in primary astrocytes, we expanded the treatment duration and analysed PMY incorporation, as well as the phosphorylation of key proteins of the m-TOR signalling cascade, a central regulatory pathway of protein synthesis. Induction of NE-mediated protein synthesis is highest at 3 h, declining at later time points, returning to basal levels after 24 h (Fig. 2A, B). The mTOR signalling pathway appears to be activated prior to the peak of protein synthesis, as seen by an increase in the levels of p-mTOR and p-p70S6K (pS6K) in the earlier treatment time points, while phosphorylation of pS6 follows the same temporal pattern of PMY incorporation (Fig. 2A, B). In line with this, inhibition of the mTOR pathway using rapamycin or torin-1 interferes with the NE-dependent induction of protein synthesis (Fig. 2C, D).

**Figure 2.**
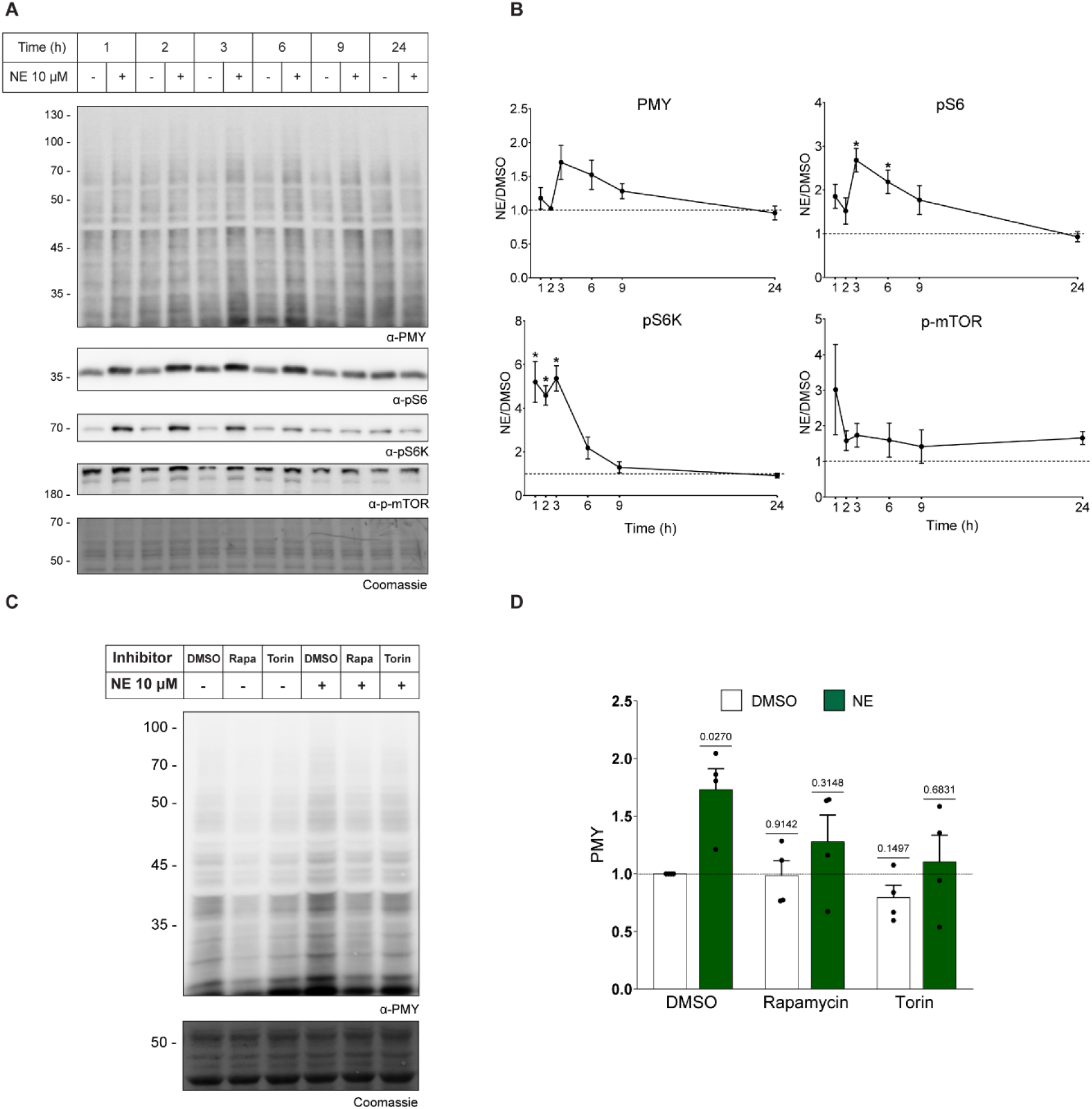
mTOR-pS6K pathway in NE-mediated protein synthesis regulation in astrocytes. **A.** Astrocytes were treated with 10 µM norepinephrine (NE, +) or DMSO (-) for 1, 2, 3, 6, 9 or 24 h and labelled with 5 µM puromycin (PMY) for 15 min immediately before sample collection. Western blot analysis was performed to detect newly synthesised proteins (anti-PMY) and phosphorylation of different elements of the mTOR pathway (S6 ribosomal protein (pS6), S6 kinase (pS6K) and mTOR (p-mTOR)). Membranes were stained with Coomassie as a loading control. **B.** Quantification of signal intensities in NE-treated samples is presented here, normalized to control treated samples for each time point. Graphs depict results from n=3 biological replicates (mean ± SEM). *p < 0.05 (one-sample t-test comparing the means with a control mean of 1). **C.** Before treatment with 10 µM NE for 3 h (or DMSO as control), astrocytes were treated with 25 nM rapamycin (Rapa),100 nM torin-1 (Torin), or DMSO for 30 min. Cells were labelled with 5 µM PMY for the final 15 min of treatment and analysed by Western blot. **D.** Quantification of PMY signal intensity normalized to DMSO-treated controls is shown as mean ± SEM for n=4 biological replicates. p-values were determined through a one-sample t-test comparing the means with a control mean of 1.

### Induction of protein synthesis by NE is mediated mainly by β-adrenergic receptors

Cortical astrocytes express a range of adrenergic receptors (ARs), so we aimed to elucidate the contribution of each subclass to the NE-mediated effect on protein synthesis. To this end, we treated cortical astrocytes with α_1_, α_2_ or β-AR agonists (phenylephrine, clonidine and isoproterenol, respectively), for up to 3 h. We observed that activating each receptor elicits a distinct temporal response in the activation of the mTOR pathway. Stimulating the α_1_- and α_2_-ARs induces a very sharp activation of S6K after 30 min of treatment, which gradually decreases over time, returning to basal levels after 2-3 h (Fig. 3, upper panel, Fig. S2). In line with this, phosphorylation of the S6 ribosomal protein also occurs at the earliest time point, and remains relatively stable for up to 2 h (Fig. 3, middle panel, Fig. S2). In contrast, stimulation of β-ARs leads to a gradual increase in the phosphorylation of both S6K and S6, reaching the higher levels of activation towards the end of the treatment (Fig. 3, upper and middle panels, Fig. S2). Interestingly, despite a pronounced activation of the mTOR pathway upon stimulation of the α_1_- and α_2_-ARs, no effect on protein synthesis (as determined by PMY incorporation) was observed (Fig. 3, lower panel, Fig. S3). On the other hand, a gradual increase in overall protein synthesis is observed following isoproterenol treatment, mirroring the temporal pattern detected upon NE treatment.

**Figure 3.**
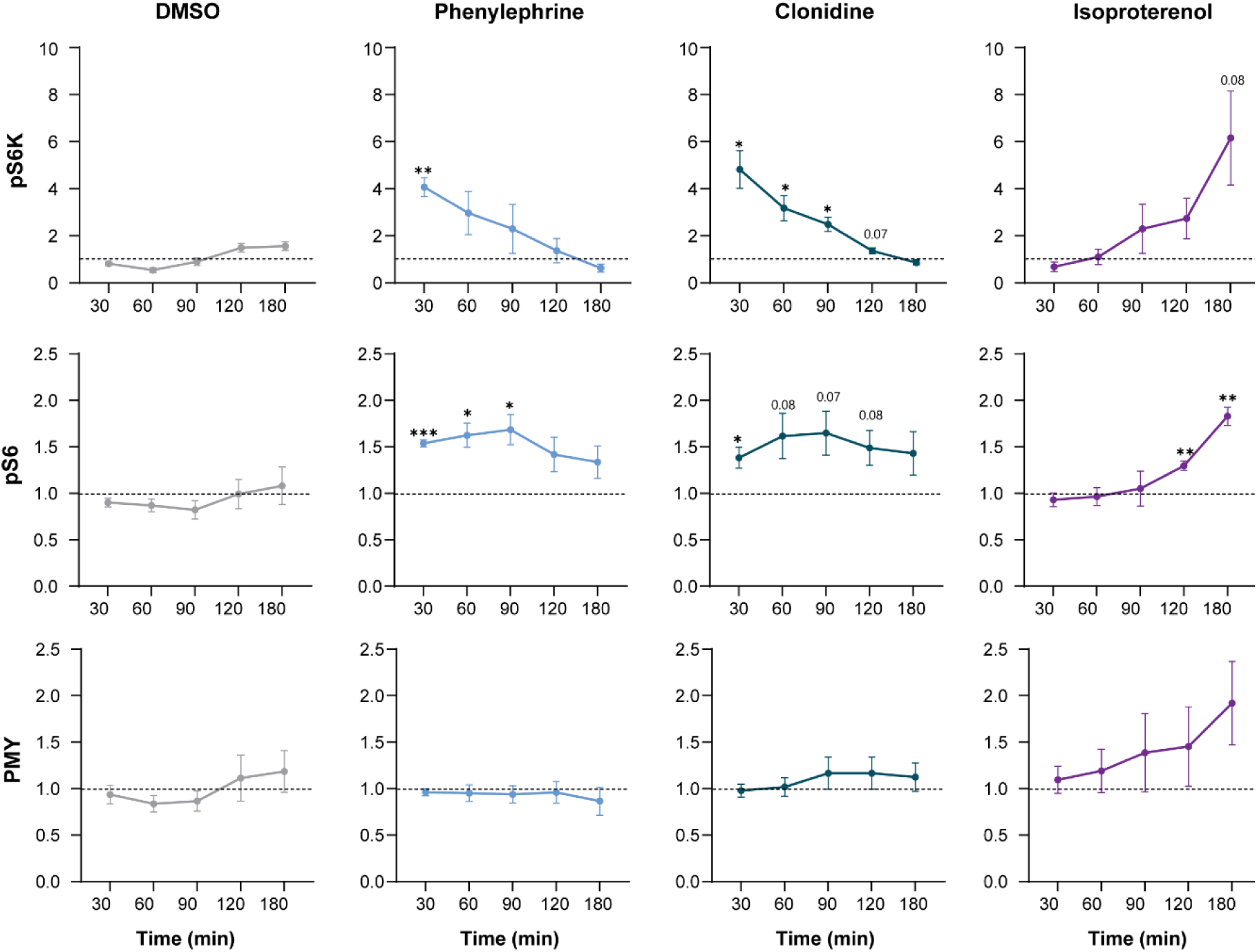
Time-dependent regulation of protein synthesis, pS6K and pS6 phosphorylation by adrenoreceptor agonists. Serum-deprived astrocytes were treated with 10 µM of the following adrenoreceptor (AR) agonists: phenylephrine (α_1_-AR), clonidine (α_2_-AR), isoproterenol (β-AR) or DMSO as a control for the indicated time points. 15 min immediately before sample collection, cells were labelled with 5 µM puromycin (PMY) and analysed by Western blot to detect phosphorylated S6 ribosomal protein kinase (pS6K, upper panel), S6 ribosomal protein (pS6, middle panel), and newly synthesised proteins (anti-PMY, lower panel). Quantification of signal intensities is presented here, normalised to untreated samples. Graphs show results from n=4 biological replicates (mean ± SEM). *p < 0.05, **p < 0.01, ***p < 0.001 (one-sample t-test comparing the means with a control mean of 1). Example blots are presented in Figure S2.

We then aimed to block each AR subtype by applying AR subtype-specific antagonists (i.e., prazosin, rauwolscine, and propranolol) either alone or combined, for 30 min prior to treatment with NE for further 3 h. Pharmacologically blocking ARs was not sufficient to prevent the effect of NE in astrocytes, as both phosphorylation of the S6 protein and overall protein synthesis were still induced in the presence of AR blockers, either on their own or in combination (Fig. S3 A, B). This suggests a complex interplay of adrenoreceptors in regulating astrocytic protein synthesis. In contrast, in C6 cells, which primarily express β-ARs (Morioka et al., 2010), pre-treatment with propranolol abolishes NE-induced protein synthesis (Fig. S3 C, D). These findings raise the possibility of crosstalk between the signalling pathways downstream of the different ARs in primary astrocytes, as well as the participation of other receptors (e.g., dopamine receptors).

β-ARs are coupled to the Gs alpha subunit, which upon activation leads to increased cAMP levels, and the activation of Protein kinase A (PKA). To determine whether a general Gs-coupled GPCR activation can induce an increase in protein synthesis, astrocytes were transduced with AAVs carrying the HA-tagged rM3D(Gs)DREADD (Guettier et al., 2009) under the GFAP promoter (Fig. S4A). Astrocytes were stimulated with CNO for 3 h and newly synthesized proteins were labelled with PMY for 15 min prior to cell collection. Stimulation of the DREADD receptor lead to a detectable increase in protein synthesis (Fig. S4B, C), suggesting that the effect involves cAMP-mediated signalling. Overall, these results indicate that the NE-dependent induction of protein synthesis occurs mainly via β-ARs.

### Protein synthesis in cultured cortical astrocytes is fuelled by glycolysis

Synthesis of new proteins is highly energetically demanding, and it has been reported that astrocytes rely mainly on glycolysis to meet their energy needs (Almeida et al., 2023). We therefore speculated that glycolysis was needed to sustain the increased rates of protein synthesis. Thus, astrocytes were pre-treated with 2-deoxy-D-glucose (2DG) for 30 min before treatment with NE for further 3 h. Inhibiting glycolysis significantly reduced protein synthesis (assessed by PMY incorporation) at basal levels, and prevented its NE-mediated induction (Fig. 4A, B), confirming that the energy required for active protein synthesis is glycolysis-derived. Of note, NE causes a decrease in astrocyte ATP levels already after a 30 min treatment, which becomes more pronounced after 1 h, showing signs of recovery at 3 h and returning to basal levels after 6 h (Fig. 4C). Such an effect is not observed upon FBS treatment, where ATP levels continue to increase from 30 min onwards (Fig. 4D).

**Figure 4.**
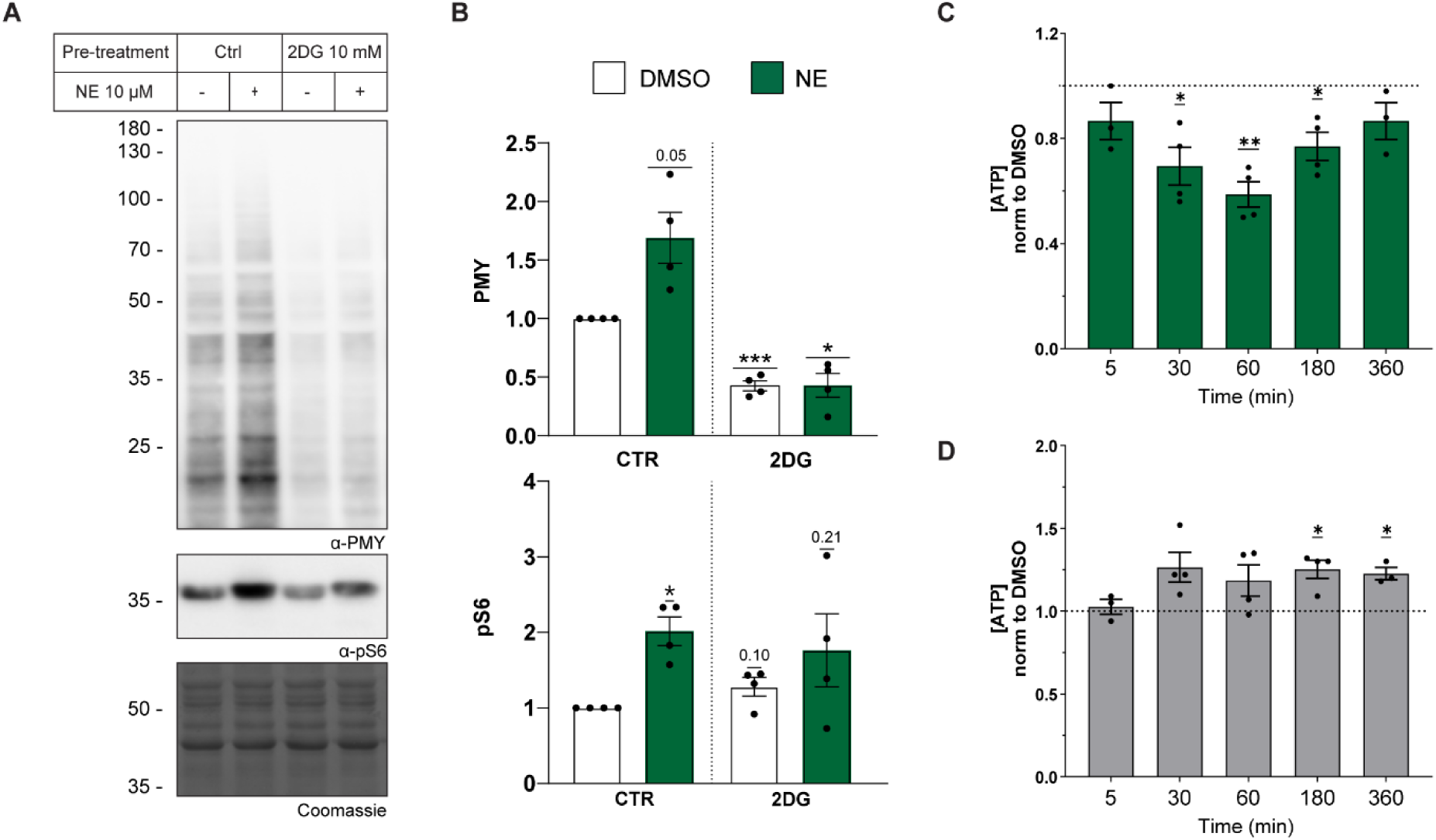
Protein synthesis in cultured astrocytes is powered by glycolysis. **A.** Astrocytes were serum-deprived in low-glucose DMEM for 24 h before pre-treatment with 10 mM 2-deoxy-D-glucose (2DG) or H_2_O as control (CTR) for 30 min, and subsequently treated with 10 µM norepinephrine (NE) or DMSO for 3 h. Cells were labelled with 5 µM puromycin (PMY) for 15 min immediately before sample collection. Samples were analysed by Western blot to detect newly synthesised proteins (anti-PMY) and phosphorylated S6 ribosomal protein (pS6). Membranes were stained with Coomassie as a loading control. **B.** Quantification of signal intensities presented here normalised to control samples (H_2_O, DMSO-treated). Graphs depict results from n=4 biological replicates (mean ± SEM). *p < 0.05, ***p < 0.001 (one-sample t-test comparing the means with a control mean of 1). **C.** Astrocytes were serum-deprived for 24 h and treated with 10 µM NE (**C**), 10% FBS (**D**) or DMSO for the indicated time points, and intracellular ATP concentrations were measured in cell lysates using a luciferase-based assay. Shown here are ATP levels in NE-(**C**) or FBS-(**D**) treated samples relative to DMSO controls, presented as mean ± SEM for n=4 biological replicates. *p < 0.05 (one sample t-test comparing the means with a control mean of 1).

## Discussion

Protein synthesis is one of the most energetically demanding processes that a cell undertakes, and as such it is tightly regulated. Efficient translational control requires the integration of information from extracellular stimuli, nutrient availability and energy status (Hershey et al., 2019; Proud, 2019). Despite its relevance, only a handful of studies have addressed the regulation of overall protein synthesis in astrocytes. An early study made use of ^35^S-methione labelling to analyse protein synthesis in mouse astrocytes at different stages of culture, as they experience either spontaneous or forskolin/dibutyryl-cAMP-induced differentiation, revealing that proteome-wide changes occur as cells undergo morphological changes (Bridoux et al., 1986). An increase in ^14^C-leucine incorporation into astrocytic proteins was reported in response to insulin (Kum et al., 1992), while 48 h cytokine treatment (TNFα and IL-1β) was shown to increase translation, as determined by ^35^S-methione/cysteine incorporation (Mandelboum et al., 2023). On the other hand, inhibition of global protein synthesis in astrocytes has been reported in response to Zn^2+^ (Alirezaei et al., 2002) or lactic acidosis (Vantelon et al., 2007). Here, we report a previously undescribed role for norepinephrine (NE), inducing overall protein synthesis in rat cortical astrocytes already apparent at ∼2 h.

Astrocytes are known to express several ARs, and NE binding to each subclass leads to the activation of distinct intracellular signalling cascades (Hertz et al., 2010; Horvat et al., 2016; Wahis and Holt, 2021). Our results show that while stimulation of all adrenoreceptor subtypes expressed in astrocytes leads to activation of the mTOR signalling pathway, an effect on protein synthesis is only detectable upon sustained application of a β-AR ligand. A more prominent role for this receptor subclass in protein synthesis stimulation is supported by our findings of a similar NE-mediated effect in C6 cells, which express a more limited repertoire of ARs, consisting mainly of β-ARs and, to a lesser extent, α_2_-ARs (Morioka et al., 2010). Additionally, we determined that general activation of Gs-coupled GPCR is sufficient to elicit an induction of protein synthesis, suggesting a role for the cAMP/PKA pathway. While our results implicate a more prominent role of the β-ARs, crosstalk between different ARs and their downstream signalling cascades has been reported (Cottingham et al., 2013), and such an interplay could also be taking place in our system, particularly in light of the inability of AR blockers to prevent an NE-mediated induction of protein synthesis. Furthermore, it is currently recognised that cross-communication extensively takes place between GPCRs and receptor tyrosine kinases (RTKs), a central class of membrane receptors, to integrate and diversify the intracellular response to signalling cues (Di Liberto et al., 2019; Kilpatrick and Hill, 2021). For instance, earlier studies have shown that stimulation of α_2_-ARs leads to the release of heparin-binding epidermal growth factor (HB-EGF) from the cell surface and the rapid phosphorylation of the EGF receptor (EGFR), resulting in ERK1/2 activation (Pierce et al., 2001). Notably, it was reported that exposure of cultured mouse astrocytes to micromolar concentrations of isoproterenol induced the shedding of growth factors, resulting in the transactivation of the EGF receptor and subsequent ERK1/2 phosphorylation via a Gs/Gi switch (Du et al., 2010). Further studies are required to better understand to what extent growth factors are released to the extracellular milieu upon NE stimulation and whether they lead to ERK and ultimately mTOR activation to enhance protein synthesis.

The protein synthesis-inducing effect of NE that we observed is somewhat delayed compared to the more rapid effects of growth-factor induced protein synthesis reported before, such as the effect of BDNF (brain-derived neurotrophic factor) or insulin on neuronal protein synthesis, readily visible after 30 min treatment (Takei et al., 2001, 2004; Liao et al., 2007). Therefore, changes in gene expression may be required to facilitate the NE-mediated effect. For instance, a recent report indicates that NE- and vasoactive intestinal peptide (VIP)-induced glycogen synthesis requires PKC- and CREB-dependent transcription (Lim et al., 2024). It remains to be determined if the activity of CREB or other transcription factors is required for the induction of protein synthesis by NE.

NE has been shown to affect astrocyte metabolism by stimulating glucose uptake, glycolysis, glycogen breakdown and lactate production. The latter is considered to be shuttled to neurons, where it is required for long-term memory formation (Alberini et al., 2018; Bak et al., 2018; Coggan et al., 2018; Zorec and Vardjan, 2023). In addition to triggering glycogen breakdown within a few minutes, NE also induces astrocyte glycogen re-synthesis in a time frame of 1-8 hours. Interestingly, this effect was mediated by β-ARs, cAMP-driven and dependent on active translation (Sorg and Magistretti, 1992; Cardinaux and Magistretti, 1996). Our results show that in addition to these effects, NE regulates protein synthesis in astrocytes, which relies on active glycolysis, thus adding another layer to the complex metabolic control NE has on astrocytes. Notably, insulin has been shown to regulate astrocytic metabolism like NE, affecting glucose, lactate and glycogen levels, as well as protein synthesis (Kum et al., 1992; Heni et al., 2011; Muhič et al., 2015).

Interestingly, within the first hours of NE treatment, we detected a decrease in total cellular ATP levels. To the best of our knowledge, this effect has not been reported before. Although protein synthesis consumes a high proportion of the cell’s energetic resources (Buttgereit and Brand, 1995; Argüello et al., 2020), our results do not support a role for the actively ongoing translation induced by NE as a cause for the observed decrease in ATP levels. Firstly, the two phenomena seem to occur at different time scales (ATP dip peaking at 60 min, protein synthesis at 3 h). Secondly, 10% FBS - a known protein synthesis-inducing stimulus - did not cause a similar transient dip. Instead of protein synthesis, other NE-triggered processes, such as cytoskeletal reorganisation and vesicle traffic may consume large amounts of ATP (Bernstein and Bamburg, 2003; Suzuki et al., 2015). Activation of β-ARs induces stellate morphology in cultured astrocytes through reorganisation of actin, microtubule and intermediate filament cytoskeletal elements (Shain et al., 1987; Goldman and Abramson, 1990; Safavi-Abbasi et al., 2001; Vardjan et al., 2014; Sherpa et al., 2016). Interestingly, morphological changes are readily visible within 30-60 min after application of β-ARs ligands (Vardjan et al., 2014), a similar time scale to the detected ATP decrease we report here.

Recent evidence has shown that NE-dependent astrocyte activation is an evolutionarily conserved mechanism of neuromodulation (Chen et al., 2025; Guttenplan et al., 2025; Lefton et al., 2025). It remains to be tested if adrenoreceptor-mediated induction of protein synthesis can be observed *in vivo* in experimental conditions eliciting elevated noradrenergic tone, and whether such a relatively slow process affects the neuromodulatory function of astrocytes. In addition, it remains to be studied what is the relevance of NE-mediated protein synthesis in the context of neurodegenerative disorders, where defects of both the noradrenergic system and proteostasis have been described (Yerbury et al., 2016; Holland et al., 2021; Zorec and Vardjan, 2023).

## Materials & Methods

### Reagents

Puromycin (PMY, 13884), norepinephrine (NE, 16673), clonidine (15949), rauwolscine (Rw, 15596), prazosin (Pz, 15023), cycloheximide (CHX, 14126), anisomycin (ANI, 11308), torin-1 (10997), rapamycin (13346) and clozapine-N-oxide (CNO, 16882) were from Cayman Chemical. Phenylephrine (HY-B0471) and isoproterenol (HY-B0468) were from MedChemExpress. DMSO (276855) and polyethylenimine (PEI, 408727) were from Sigma. 2-deoxy-D-glucose (2DG, 11980050) was from Thermo Scientific and propranolol (Prop, 0624) was from Tocris Bioscience.

### Cell culture & treatments

Primary cortical astrocytes were isolated from embryonic day 21 (E21) Sprague Dawley rat pups, using a modified McCarthy and DeVellis method (McCarthy and de Vellis, 1980). Cerebral cortices were digested with 0.25% Trypsin-EDTA (Invitrogen) for 20 min at 37°C, with addition of 0.5 mg/mL DNAse I (Roche Diagnostics) and 10 mM MgSO_4_ after the first 10 min. Digestion was terminated by addition of 0.25% trypsin inhibitor (Invitrogen), and 1% BSA, and tissue was mechanically disrupted with 1mL pipette tips. Tissue suspension was then diluted in HBSS, large debris removed by 30 sec centrifugation at 200g, and cells collected by 5 min centrifugation at 200g. Cells were then seeded on poly-L-lysine (PLL)-coated 75 cm^2^ flasks, and maintained in Dubelcco’s Modified Eagle Medium (DMEM, Gibco) supplemented with 10% foetal bovine serum (FBS, Gibco, Performance Plus), 100 U/mL penicillin and 100 μg/mL streptomycin (Invitrogen), with medium changes every 48 h. On DIV 6, medium was supplemented with 10 mM HEPES (pH 7.4) and flasks were shaken at 180 rpm on a 50 mm orbital throw shaker at 37°C, overnight. After, medium was removed, cells were washed twice with PBS and fresh medium was added. On DIV 8, cells were detached from the flask with 0.08% Trypsin-EDTA and plated in PLL-coated plates for subsequent experiments. Cortical neurons were also prepared from E21 rat pup cortices, following a similar tissue digestion and suspension procedure, but maintained in Neurobasal A medium (Invitrogen), supplemented with B27 (Invitrogen), 1mM L-glutamine (Invitrogen) and 100 µg/mL Primocin (InvivoGen), in the presence of 10 µM FdU, with half-medium changes every 48 h. C6 glioma and HMC3 microglial cells were cultured in DMEM supplemented with 10% FBS (Pan Biotech), 100 U/mL penicillin and 100 μg/mL streptomycin (Invitrogen). All cells were cultured in a humidified incubator at 37°C with a 5% CO_2_ atmosphere. Relevant procedures were approved by the ethics committee of animal experiments at the Ministry of Agriculture of Estonia (Permit Number: 45).

Prior to pharmacological treatments, cells were serum-deprived (or B27-deprived in the case of neurons) for 24 h, and subsequently treated with either 2% B27, 10% FBS, norepinephrine (NE), clonidine, phenylephrine or isoproterenol, all 10 µM. When indicated, cells were pre-treated with either torin-1 (100 nM), rapamycin (25 nM), anisomycin (10 µg/mL), prazosin, rauwolscine or propranolol (all 10 µM) for 30 min before addition of NE. For treatment with 10 mM 2-deoxy-D-glucose (2DG), astrocytes were serum-deprived for 24 h in glucose-free DMEM (Gibco, 11966025), supplemented with 1 mg/mL glucose (Sigma, G-8270). As control, cells were treated with 0.1% DMSO, whenever indicated. For puromycin (PMY) labelling, cells were treated with 5 µM PMY for 15 min before lysis or 2.5 min before fixation for ICC.

### Western blotting

Equal amounts of lysates were separated by Tris-Glycine SDS-PAGE following Laemmli’s discontinuous buffer system and transferred to PVDF membranes using Trans-Blot Turbo Transfer system (Bio-Rad). The following antibodies were used for immunoblot: anti-puromycin (Millipore MABE343, 1:2000), anti-phospho-S6 Ribosomal Protein (Ser240/244) (Cell Signaling, D68F8, 1:2000), anti-phospho-p70 S6 Kinase (Thr389) (Cell Signaling, 108D2, 1:1000) and anti-phospho-mTOR (Ser2448) (Cell Signaling, D9C2, 1:1000). HRP-conjugated secondary antibodies (anti-mouse IgG, 32430 and anti-rabbit IgG, 32460, Thermo Scientific™) were used at a 1:5000 dilution. Immunoblots were developed with SuperSignal™ West Dura/Atto Maximum Sensitivity Substrate (Thermo Scientific™) and images captured with ImageQuant LAS 4000 imaging system (GE Healthcare Life Sciences). For equal loading control, membranes were stained with Coomassie. Signal intensity quantification was performed with the IOCBIO Gel Software (Kütt et al., 2023).

### Immunocytochemistry (ICC)

Following treatment and labelling, astrocytes grown on coverslips were fixed with 4% paraformaldehyde-PBS for 15 min, permeabilized with 0.5% Triton X-100 in PBS for 15 min and blocked with 2% BSA-PBS for 1 h. Incubation with primary antibodies was performed overnight at 4°C, in 0.2% BSA-PBS, using the following antibodies: anti-puromycin (Millipore MABE343, 1:2000), anti-GS (Sigma, G2781, 1:2000), anti-HA (Cell Signaling, C29F4, 1:1000) and anti-Aldh1L1 (DSHB, N103/31, 1:200). After washing, incubation with fluorophore-conjugated secondary antibodies (anti-mouse or anti-rabbit Alexa Fluor-594 and Alexa Fluor-488, Thermo Scientific) was done for 1-2 h at RT, in 0.2% BSA-0.1% in the dark. Lastly, cells were counterstained with Hoechst 33342 (Thermo Scientific, 5 µg/mL) before mounting in ProLong Gold antifade mounting medium (Thermo Scientific). Images were obtained using a Zeiss LSM 900 confocal microscope and puromycin signal intensity was quantified under the GS-positive cells using ImageJ. 10 images per condition and biological replicate were obtained, the puromycin integrated density was determined under a GS mask and normalized by the measured area.

### Metabolic labelling with ^35^S methionine/cysteine

Astrocytes were grown in 12-well plates, serum-deprived for 24 h in DMEM lacking methionine and cysteine (Gibco, 21013-024) and supplemented with 4 mM glutamine, treated with 10 µM NE for 3 h (or DMSO as control) and pulse labelled for 1 h at the end of the treatment with 25 µCi/mL ^35^S methionine/cysteine (EasyTag Express NEG772002MC, Revvity). 100 µg/mL cycloheximide was added together with the ^35^S methionine/cysteine label where indicated. At the end of incubation, cells were gently washed twice with 1 mL PBS at room temperature, lysed directly in 150 µL Laemmli buffer and boiled. 50 µL of these lysates were combined with 10 µL of 20 mg/mL bovine serum albumin and total protein was precipitated for 15 min on ice by equal volume (60 µL) of ice cold 20% TCA. Precipitates were pelleted by centrifugation for 10 min at 16000 g at 4°C, washed once with 0.5 mL ice cold acetone and resuspended in 100 µL 1M NaOH. Protein samples were mixed with 3 mL Ultima Gold scintillation cocktail and analysed using the Quantulus 1220 scintillation counter (Perkin Elmer).

### Intracellular ATP measurements

ATP levels were determined using a luciferase-based assay (ATP Detection Assay Kit – Luminescence, Cayman Chemical, 700410). For this, astrocytes were plated onto 48-well plates and grown to confluency before serum-deprivation for 24 h. Cells were then treated with 10 µM NE, 10% FBS or 0.1% DMSO for 5, 30, 60, 120, 180 and 360 min. Following treatment, medium was removed, cells washed with ice-cold PBS and lysed following the manufacturer’s instructions. Luminescence signal was measured using a Genios Pro Multifunction Microplate Reader (Tecan), in 96-well plates. ATP concentrations were determined against an ATP standard curve, as specified in the user manual. ATP levels were normalized to those of DMSO-treated cells for each experimental replicate.

### Adeno-associated virus (AAV) production, purification and transduction

AAVs were generated using the AAV-293 cell line (Agilent, 240073), grown in DMEM supplemented with 10% FBS,100 U/mL penicillin and 100 mg/mL streptomycin and transfected at 60-70% confluency with PEI (2:1 weight ratio of PEI and DNA). Four 10 cm plates were transfected with 10 μg of DNA each, consisting of: pAAV-GFAP-HA-rM3D(Gs)-IRES-mCitrine plasmid (Addgene, Plasmid #50472), pAdDeltaF6 helper plasmid, AAV1 and AAV2 helper plasmids (Addgene, Plasmid #112867, Plasmid #112862, and Plasmid #104963, respectively) at 1:1:0.5:0.5 molar ratios. 72 h after transfection virus particles were collected and purified using AAVpro® Purification Midi Kit for All Serotypes from Takara Bio (#6675). Virus titers (as determined by RT-qPCR) were in the range of 10^11^ viral genomes/mL. For infection, astrocytes were detached from the growth flasks (1-2 days after shaking) and incubated with AAVs (10^10^ viral genomes/mL) at the time of seeding onto cell culture plates. Infection was visually confirmed via expression of the mCitrine tag. 6 days after infection, cells were serum-deprived for 24 h and subsequently stimulated with 5 µM CNO or 0.1% DMSO as control for 3 h.

## Author Contributions

The study was conceived by R.Z., I.K. and F.C-C. Experiments were designed and supervised by F.C.-C. and I.K. Experiments were carried out by F.C-C., H.T., G.U, and A.U. The manuscript was written by F.C.-C. and I.K., and revised by all authors.

## Acknowledgments

This work was supported by the Estonian Research Council (Grants PRG2206 to I.K., MOB3JD1210 to F.C.C). We thank Epp Väli for technical support, and Annela Avarlaid, Olga Jasnovidova and Tõnis Timmusk for insightful discussions. RZ is acknowledging the support by the Slovenian Research and Innovation Agency grants: P3-0310, J3-2523, J3-50104, MR+, I0-0034, I0-0022), MRIC-Carl Zeiss Reference Centre for Laser Confocal Microscopy (P3-0083), MRIC UL (ICBSL3+; grant I0-0510), COST action (CA18133; ERNEST).

