## Supplementary material for "Norepinephrine stimulates protein synthesis in astrocytes": Supplemetary Figures

### Supplementary figures

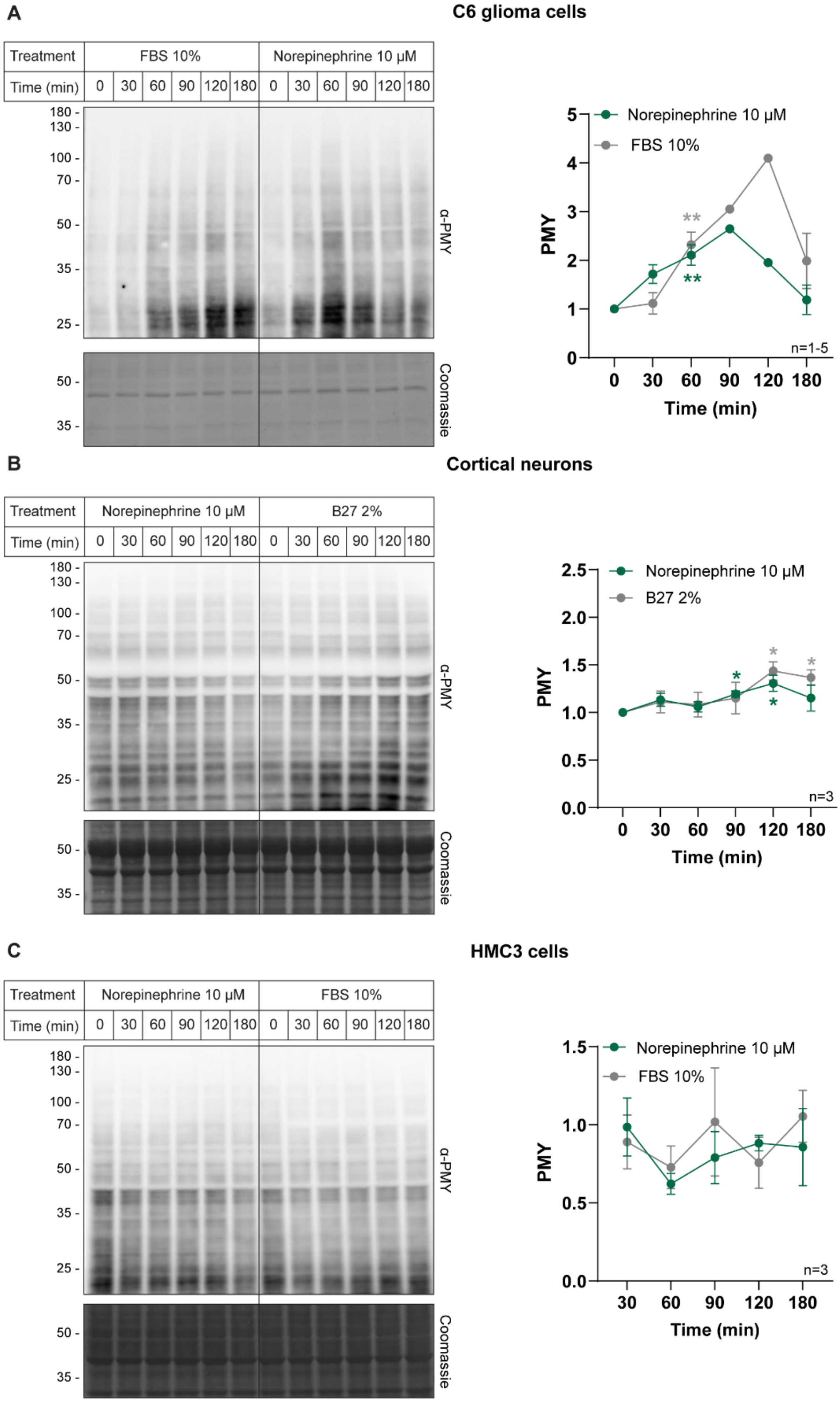

**Figure S1. Effect of norepinephrine on protein synthesis in other neural cells.**

C6 glioma cells (**A**), primary cortical neurons (**B**) or HMC3 microglial cells (**C**), were serum- or B27-deprived for 24 h and subsequently treated with 10 μM norepinephrine (NE), 2% B27 or 10% FBS, before labelling with 5 μM puromycin (PMY) for 15 min immediately before harvesting. Samples were analysed by Western blot to detect newly synthesised proteins (anti-PMY). Membranes were stained with Coomassie as a loading control. Quantification of signal intensities is presented here, normalised to control samples (DMSO). Graphs depict mean ± SEM. $^{*}p < 0.05$, $^{**}p < 0.01$ (one-sample t-test comparing the means with a control mean of 1). The number of biological replicates is shown next to the quantification in each panel.

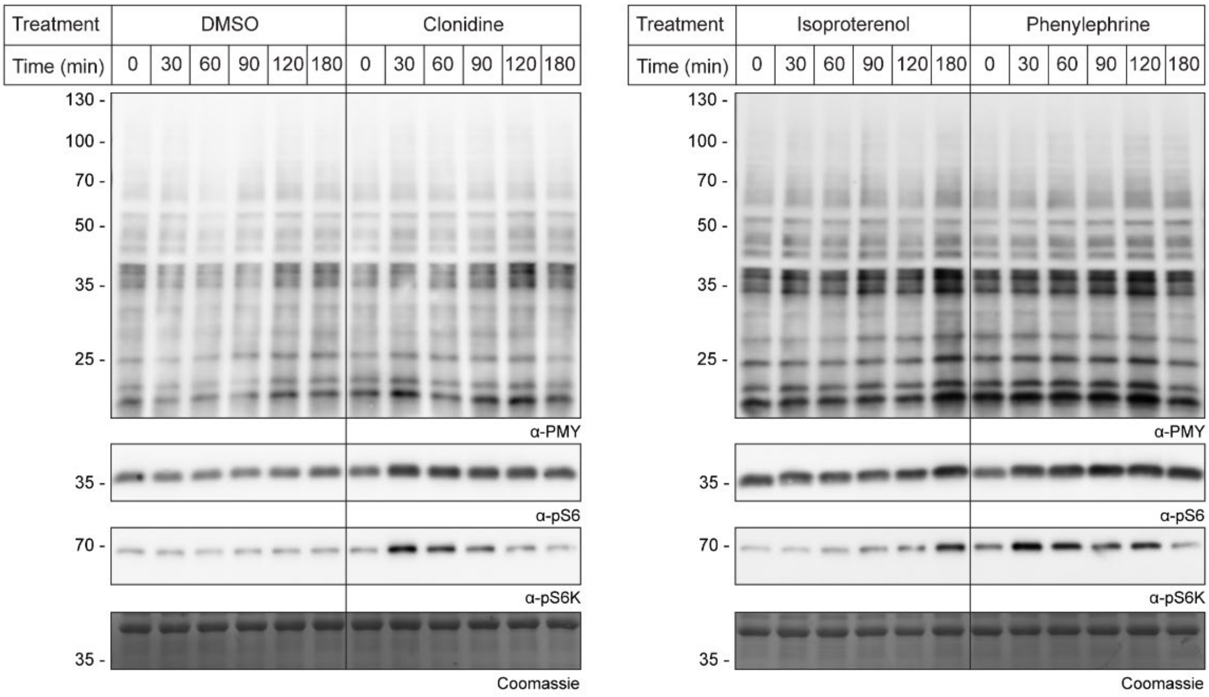


**Figure S2. Time-dependent regulation of protein synthesis, pS6K and pS6 phosphorylation by adrenoreceptor agonists (related to Figure 3).**

Example blots related to Figure 3. Astrocytes were serum-deprived for 24 h and then treated with 10 μM of either of the following adrenoreceptor (AR) agonists: phenylephrine ($\alpha_1$-AR), clonidine ($\alpha_2$-AR), isoproterenol (β-AR) or DMSO as a control for the indicated time points. Cells were labelled with 5 μM puromycin (PMY) 15 min immediately before sample collection and analysed by Western blot to detect newly synthesised proteins (anti-PMY), phosphorylated ribosomal protein (pS6) and phosphorylated S6 kinase (pS6K). Membranes were stained with Coomassie as a loading control.

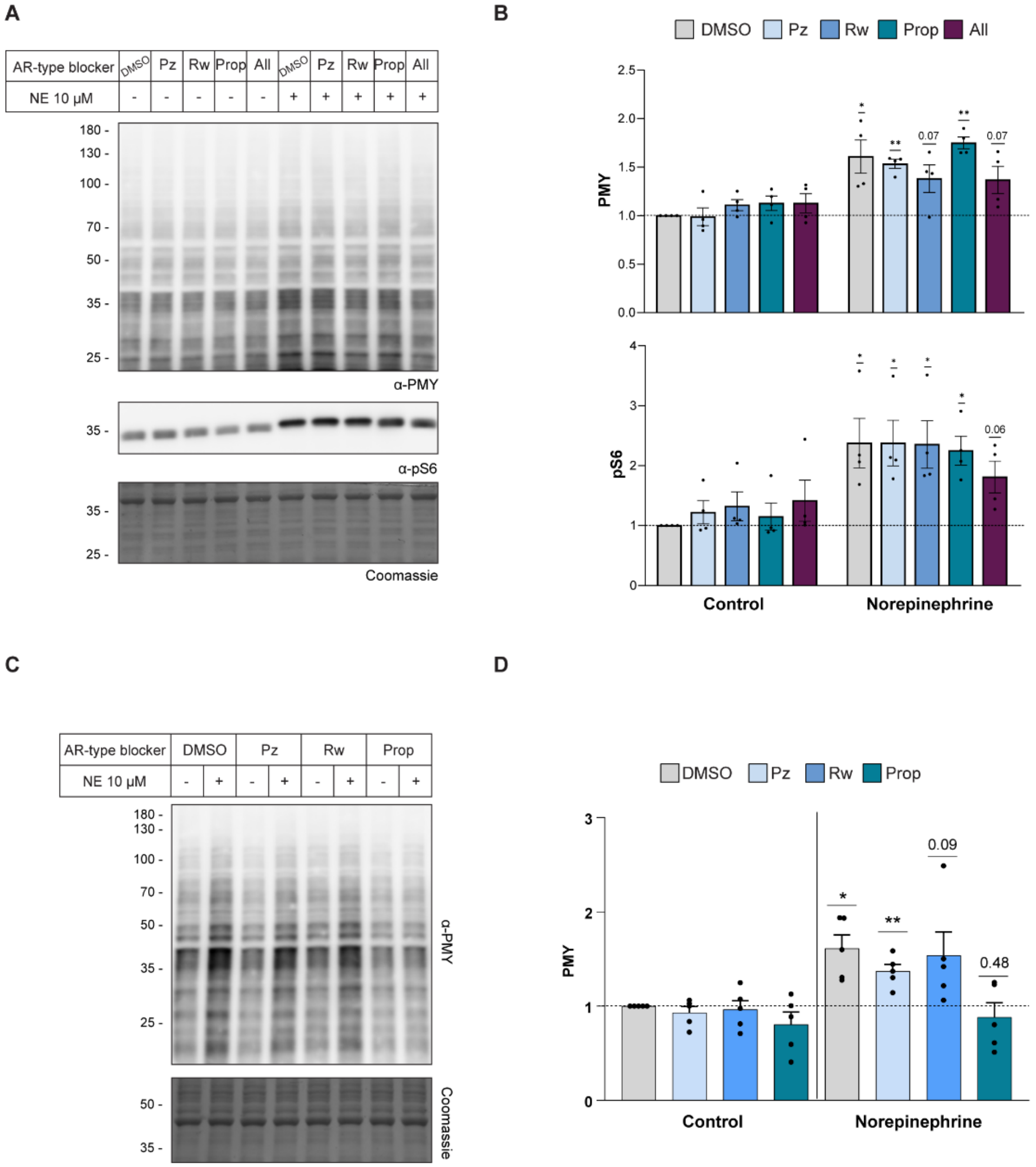


**Figure S3. The effect of selective adrenergic antagonists on protein synthesis in primary astrocytes and C6 glioma cells.**

**A.** Serum-deprived primary cultures of rat cortical astrocytes were pre-treated with one of the following adrenergic receptor (AR) blockers: prazosin (Pz), rauwolscine (Rw), propranolol (Prop), or a combination of the three (All); all 10 µM. Control cells were pre-treated with DMSO. After 30 min, cells were treated with 10 µM norepinephrine (NE) or DMSO as control, for 3 h, and newly synthesised proteins were labelled with 5 µM puromycin (PMY) during the final 15 minutes of treatment. Samples were analysed by Western Blot using anti-PMY and antiphosphorylated S6 ribosomal protein (pS6) antibodies. Membranes were stained with Coomassie as a loading control.

**B.** Quantification of signal intensity normalized to DMSO controls is shown as mean ± SEM for n=4 biological replicates. *p < 0.05, **p < 0.01 (one sample t-test comparing the means with a control mean of 1).

**C.** C6 glioma cells were serum-deprived for 24 h and pre-treated with the indicated AR blockers for 30 min (same concentrations as indicated above), before incubation with 10 µM NE (or DMSO as control) for 2 h. Newly synthesised proteins were labelled with 5 µM PMY for 15 min and analysed by Western blot using an anti-PMY antibody. Coomassie staining was included as a loading control.

**D.** Signal intensities were quantified from n=5 biological replicates and presented here relative to control samples (DMSO-treated). Graph shows mean ± SEM. *p < 0.05, **p < 0.01 (one sample t-test comparing the means with a control mean of 1).

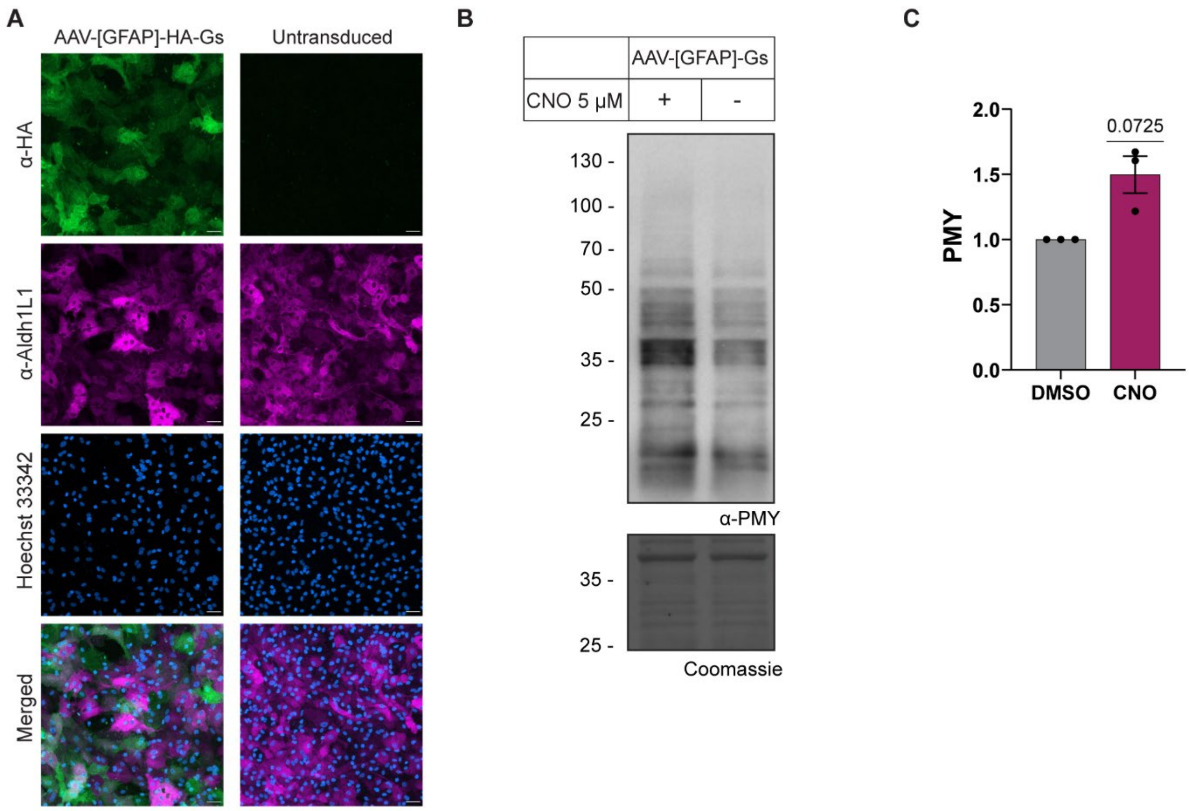


**Figure S4. Gs-DREADDs-mediated activation of protein synthesis in astrocytes.**

**A.** Astrocytes transduced with AAVs carrying an HA-tagged rM3D(Gs)DREADD under the GFAP promoter ("AAV-[GFAP]-HA-Gs") were fixed 7 days after infection and analysed by immunocytochemistry using an anti-HA antibody to visualize the expression of the Gs-DREADD construct (shown in green) and an anti-Aldh1L1 antibody as an astrocyte marker (shown in magenta). Scale bar: 50 μm.

**B.** Transduced astrocytes were serum-deprived for 24 h and subsequently treated with 5 μM of the DREADD-specific ligand clozapine-N-oxide (CNO) or DMSO as control for 3 h. Cells were labelled with 5 μM puromycin (PMY) for 15 min immediately before collection. Samples were analysed by Western blot to detect newly synthesised proteins (anti-PMY). Membranes were stained with Coomassie as a loading control.

**C.** Quantification of PMY signal intensity is presented here normalized to DMSO-treated samples. Graphs depict results from n=3 biological replicates as mean ± SEM; one sample t-test comparing the means with a control mean of 1.
